# Non-linear manifold learning in fMRI uncovers a low-dimensional space of brain dynamics

**DOI:** 10.1101/2020.11.25.398693

**Authors:** Siyuan Gao, Gal Mishne, Dustin Scheinost

**Affiliations:** Department of Biomedical Engineering, Yale University, United States, 06520; Halıcıoğlu Data Science Institute, University of California San Diego, United States, 92093; Neurosciences Graduate Program, University of California San Diego, United States, 92093; Department of Radiology and Biomedical Imaging, Yale School of Medicine, United States, 06510; Department of Statistics and Data Science, Yale University, United States, 06520; Child Study Center, Yale School of Medicine, United States, 06510

## Abstract

Large-scale brain dynamics are believed to lie in a latent, low-dimensional space. Typically, the embeddings of brain scans are derived independently from different cognitive tasks or resting-state data, ignoring a potentially large—and shared—portion of this space. Here, we establish that a shared, robust, and interpretable low-dimensional space of brain dynamics can be recovered from a rich repertoire of task based fMRI data. This occurs when relying on non-linear approaches as opposed to traditional linear methods. The embedding maintains proper temporal progression of the tasks, revealing brain states and the dynamics of network integration. We demonstrate that resting-state data embeds fully onto the same task embedding, indicating similar brain states are present in both task and resting-state data. Our findings suggest analysis of fMRI data from multiple cognitive tasks in a low-dimensional space is possible and desirable, and our proposed framework can thus provide an interpretable framework to investigate brain dynamics in the low-dimensional space.

## Introduction

Understanding large-scale brain dynamics is a major goal of modern neuroscience (Jorgenson et al., 2015). However, due to the high-dimensional nature of brain patterns, how to best operationalize and tackle this problem remains an open question. Nevertheless, the temporal dimensions that explain the observed dynamics is small compared with the number of time points (Cunningham and Byron, 2014). Thus, there is growing evidence to suggest that a low dimensional space—hidden from direct observation, learned from the data, and derived from many brain regions—may be a suitable model for studying temporal brain dynamics (Gao and Ganguli, 2015).

These low-dimensional spaces have been observed using a variety of neural recordings and animal models (Ahrens et al., 2012; Churchland et al., 2012; Kobak et al., 2016; Mishne et al., 2016; Santhanam et al., 2009). Research suggests that linear methods, such as principal component analysis (PCA), are appropriate when recorded temporal data comes from simple stimuli that project onto a limited area within a manifold (Cunningham and Byron, 2014). However, data from richer tasks often project onto a larger portion of the manifold, violating linear approximations (Cunningham and Byron, 2014; Gallego et al., 2017). Nonlinear dimensionality reduction methods, like diffusion maps (Coifman and Lafon, 2006), can overcome this limitation by integrating local similarities into a global representation, which better reflect the underlying temporal dynamics in neural recordings.

Similar concepts have emerged in human functional magnetic resonance imaging (fMRI) studies to quantify moment-to-moment changes in activity and connectivity (Hutchison et al., 2013b; Preti et al., 2017). As with related research on temporal recordings from animal models, dimensionality reduction methods are used to project the fMRI time series onto a low-dimensional space (Allen et al., 2014b; Monti et al., 2017; Shine et al., 2016; Shine et al., 2019). From the low-dimensional space, characteristic brain states—or distinct, repeatable patterns of brain activity—are used to quantify brain dynamics. Predominantly, these studies have relied on linear methods (Allen et al., 2014b; Monti et al., 2017; Shine et al., 2016; Shine et al., 2019). However, given the rich repertoire of tasks available in human fMRI, a manifold derived from nonlinear methods may better capture the underlying geometry of the low-dimensional space.

To address this, we recently introduced 2-step Diffusion Maps (2sDM; Gao et al., 2019), which is a novel extension of diffusion maps. 2sDM extracts common variability between individuals by performing dimensionality reduction of a 3rd-order tensor in a two-stage manner. In the first stage, timeseries data from each individual are embedded into a low-dimensional Euclidean space. In the second stage, embedding coordinates for the same time point from different individuals are concatenated for use in a second embedding. The second stage embeds similar time points across subjects to obtain a low-dimensional group-wise representation of those time points. This two-stage approach avoids directly comparing brain activation across subjects, which can be imprecise without proper alignment (Haxby et al., 2011). As 2sDM is an unsupervised learning method, there is no need to handcraft features, which are less robust, computationally intensive, and generalize poorly when compared to learned features from unsupervised methods (Bengio et al., 2013). While diffusion maps have been applied to fMRI data (Margulies et al., 2016; Nenning et al., 2020), we aim to embed the time dimension rather than the spatial dimension.

We used 2sDM to embed timeseries from a rich repertoire of tasks onto a single low-dimensional manifold in two fMRI datasets: the Human Connectome Project and the UCLA Consortium for Neuropsychiatric Phenomics. By using multiple tasks spanning a range of cognitive functions and loads, we obtain a more even sampling of the original high-dimensional space of recurring patterns of brain dynamics (Cunningham and Byron, 2014; Gallego et al., 2017) to better project individual time points onto a low-dimensional manifold. Compared with other embedding results that only aim to separate different tasks, our embedding positioned different tasks by their cognitive load. Thus, it enables scans to be compared both within the same task and across different tasks. As our embedding also has a clear clustering structure, downstream analyses that are based on brain states or low-dimensional trajectories are also straightforward to perform based on the embedding. Additionally, we embedded resting-state data into the same task embedding to investigate differences in brain dynamics between resting-state and task performance. These results suggest that manifold learning can uncover an interpretable low-dimensional embedding for the study of brain dynamics in fMRI data.

## Methods

### Dataset and imaging parameters

Data was obtained from the Human Connectome Project (HCP) 900 Subject release (Van Essen et al., 2013). We use fMRI data collected while 390 participants performed six tasks (gambling, motor, relational, social, working memory—WM, and emotion). We restrict our analyses to those subjects who participated in all nine fMRI conditions (seven task, two rest), whose mean frame-to-frame displacement is less than 0.1mm and whose maximum frame-to-frame displacement is less than 0.15mm, and for whom the task block order is the same as other subjects (*n* = 390). All fMRI data were acquired on a 3T Siemens Skyra using a slice-accelerated, multiband, gradient-eco, echo planar imaging (EPI) sequence (TR=720ms, TE=33.1ms, flip angle=52°, resolution=2.0mm^3^, multiband factor=8). Images acquired for each subject include a structural scan and eighteen fMRI scans (working memory (WM) task, incentive processing (gambling) task, motor task, language processing task, social cognition task, relational processing task, emotion processing task, and two resting-state scans; two runs per condition (one LR phase encoding and one RL phase encoding run)) split between two sessions.

The UCLA Consortium for Neuropsychiatric Phenomics (CNP; Poldrack et al., 2016) dataset is used for replication. Similar to the standards for the HCP dataset, we restrict our analyses to those subjects who participated in all 5 fMRI conditions (four task, one rest), whose mean frame-to-frame displacement is less than 0.1mm and whose maximum frame-to-frame displacement is less than 0.15mm. 77 healthy controls are retained. These participants performed four tasks (paired memory retrieval task—PAMRET, paired memory encoding task—PAMENC, spatial working memory task—SCAP, task switching task—TASKSWITCH). Details of the image acquisition parameters have been published elsewhere (Poldrack et al., 2016). In brief, all data were acquired on one of two 3T Siemens Trio scanners at UCLA. Functional MRI data were collected using a T2*-weighted EPI sequence with the following parameters: slice thickness=4mm, 34 slices, TR=2s, TE=30ms, flip angle=90°, matrix 64×64, FOV=192mm, oblique slice orientation. Images acquired for each subject include a structural scan and seven fMRI scans (balloon analog risk task (BART), paired-associate memory retrieval (PAMRET), paired-associate memory encoding (PAMENC), spatial capacity task (SCAP), stop signal task (SST), task-switching task (TASKSWITCH) and breath holding task).

As 2sDM requires time series to be synchronized across individuals (i.e., different individuals encounter the same task condition at the same time point), the language task from the HCP and the stop signal task, balloon analogue risk task, and breath hold task from the CNP were not included. These tasks are self-paced. Participants finished blocks at different times, causing the task block to be unsynchronized across participants.

### fMRI processing

For the HCP dataset, the HCP minimal preprocessing pipeline was used (Glasser et al., 2013), which includes artifact removal, motion correction, and registration to standard space. For the CNP dataset, structural scans were skull-stripped using OptiBet (Lutkenhoff et al., 2014) and registered to the MNI template using a validated algorithm in BioImage Suite (Joshi et al., 2011; Scheinost et al., 2017). Slice time and motion correction were performed in SPM8. For both datasets, all subsequent preprocessing was performed using image analysis tools available in BioImage Suite and included standard preprocessing procedures (Finn et al., 2015). Several covariates of no interest were regressed from the data including linear and quadratic drifts, mean cerebral-spinal-fluid (CSF) signal, mean white-matter signal, and mean gray matter signal. For additional control of possible motion related confounds, a 24-parameter motion model (including six rigid-body motion parameters, six temporal derivatives, and these terms squared) was regressed from the data. The data were temporally smoothed with a Gaussian filter (approximate cutoff frequency=0.12Hz). Mean frame-to-frame displacement yielded seven motion values per subject, which were used for subject exclusion and motion analyses. We restricted our analyses to subjects whose maximum frame-to-frame displacement was less than 0.15mm and mean frame-to-frame displacement was less than 0.1mm. This conservative threshold for exclusion due to motion was used to mitigate the effect of motion on the embedding. We used the Shen 268-node atlas to extract timeseries from the fMRI data for further analysis (Shen et al., 2013). Timeseries used for the embedding were the average of the basis of the “raw” task time courses, with no removal of task-evoked activity, for each node in the atlas. Finally, 2sDM was applied to embed a 3rd-order tensor of fMRI data (individual × region × time) onto a single low-dimensional manifold.

### 2-step diffusion maps (2sDM)

Diffusion maps are part of a broad class of manifold learning algorithms. Specifically, diffusion maps provide a global description of the data by considering only local similarities and are robust to noise perturbations. The new nonlinear representation provided by diffusion maps reveals underlying intrinsic parameters governing the data (Coifman and Lafon, 2006). We briefly describe the diffusion maps algorithm in general and in the following its application to fMRI data as used in our approach. Given a dataset of n points 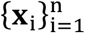 a pairwise similarity matrix **S** between pairs of data points **x**_*i*_ and **x**_*j*_ is constructed, for example using the Gaussian kernel *w*_ϵ_ (**x**_i_, **x**_*j*_)= exp (−∥**x**_i_ – **x**_*j*_∥^2^/ϵ). Then the rows of the similarity matrix are normalized by **P** = **D**^−**1**^**S**, where **D**_*ii*_ = ∑_*j*_ **S**_*ij*_ is the degree of point **x**_i_. This creates a random walk matrix on the data with entries set to *p*(**x**_*i*_, **x**_*j*_)= *w*_ϵ(_**x**_i_, **x**_j)_/*d*(**x**)_*i*_. Taking the t-th powers of the matrix **P** is equivalent to running the Markov chain corresponding to the random walk on the data forward t times. The corresponding kernel *p*_t_(∙,∙) can then be interpreted as the transition probability between two points in t time steps. The matrix **P** has a complete sequence of bi-orthogonal left and right eigenvectors ϕ_*i*_ and ψ_*i*_, respectively, and a corresponding sequence of eigenvalues 1 = 03BB_0_ ≥ |λ_1_| ≥ |λ_2_| ≥ ⋯. Diffusion maps are a nonlinear embedding of the data points into a low-dimensional space, where the mapping of point **x** is defined as 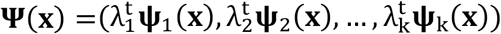, where t is the diffusion time. Note that ψ_0_ is neglected because it is a constant vector. The diffusion distance 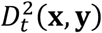 between two data points is defined as:

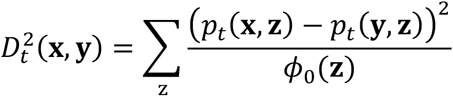

where ϕ_0_ represents the stationary distribution of the random walk described by the random walk matrix **P**. This measures the similarity of two points by the evolution in the Markov chain and the distance characterizes the probability of transition from **x** or **y** to the same **z** point in t time steps. Two points are closer with smaller 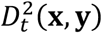 if there is a large probability of transition from **x** to y or vice versa, suggesting that there are more short paths connecting them. It is thus robust to noise as it considers all the possible paths between two points and is thus less sensitive to noisy connections. It was proved that the *k*-dimensional diffusion maps ψ embed data points into a Euclidean space ℝ^k^ where the Euclidean distance approximates the diffusion distance (Coifman and Lafon, 2006). In practice, eigenvalues of **P** typically exhibit a spectral gap such that the first few eigenvalues are close to one with all additional eigenvalues being much smaller than one. Thus, the diffusion distance can be well approximated by only the first few eigenvectors. Therefore, we can obtain a low-dimensional representation of the data by considering only the first few eigenvectors of the diffusion maps. Intuitively, diffusion maps embed data points closer when it is harder for the points to escape their local neighborhood within time t.

To obtain a groupwise low-dimensional representation of dynamics, a hierarchical diffusion maps-based manifold learning framework, 2-step diffusion maps (2sDM; Gao et al., 2019), was used to reduce the dimensionality of high-dimensional multi-individual fMRI time series. The framework is illustrated in Figure 1a. Under the assumption that individuals’ fMRI responses are time-synchronized, the fMRI BOLD time series data are organized as three-dimensional array **X** ∈ ℝ^*M*×*V*×*T*^ (number of individuals *M*, number of regions or voxels *V*, number of time points *T*). In the first step of 2sDM, each individual is processed separately, by applying diffusion maps to the fMRI time series of every single individual **X**_*i⋅*_ ∈ ℝ^*V*×*R*^, thereby obtaining a *d*_1_-dimensional temporal embedding of each individual 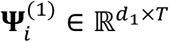. Then, in the second step, we first concatenate the new representations from all individuals into a matrix 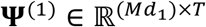, such that each time-point is now represented by the embeddings of that time-frame aggregated from all *M* subjects. Then, a second-round diffusion embedding is performed, further reducing the dimensionality of every time-frame to *d*_’_ and the final time-frame representation with multi-individual similarity is 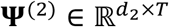, where *d*_1_ and *d*_2’_ are predetermined parameters that are smaller than *V*. The concatenation and two-round embeddings are theoretically supported by the theorem that low-dimensional diffusion maps approximate the diffusion distance between time-frames (Gao et al., 2019): The distance between two frames ψ^(2)^(t_*i*_) and ψ^(2)^(t_*j*_) approximates the average diffusion distance between those time-frames across all individuals. We used *d*_1_ = 7 and *d*_*2*’_ = 3 in our experiment. It is worth noting that the embedding results were robust under a certain range of different *d*_1_ and *d*_2_ (related discussion in Supplementary Materials and Figure S3).

**Figure 1.**
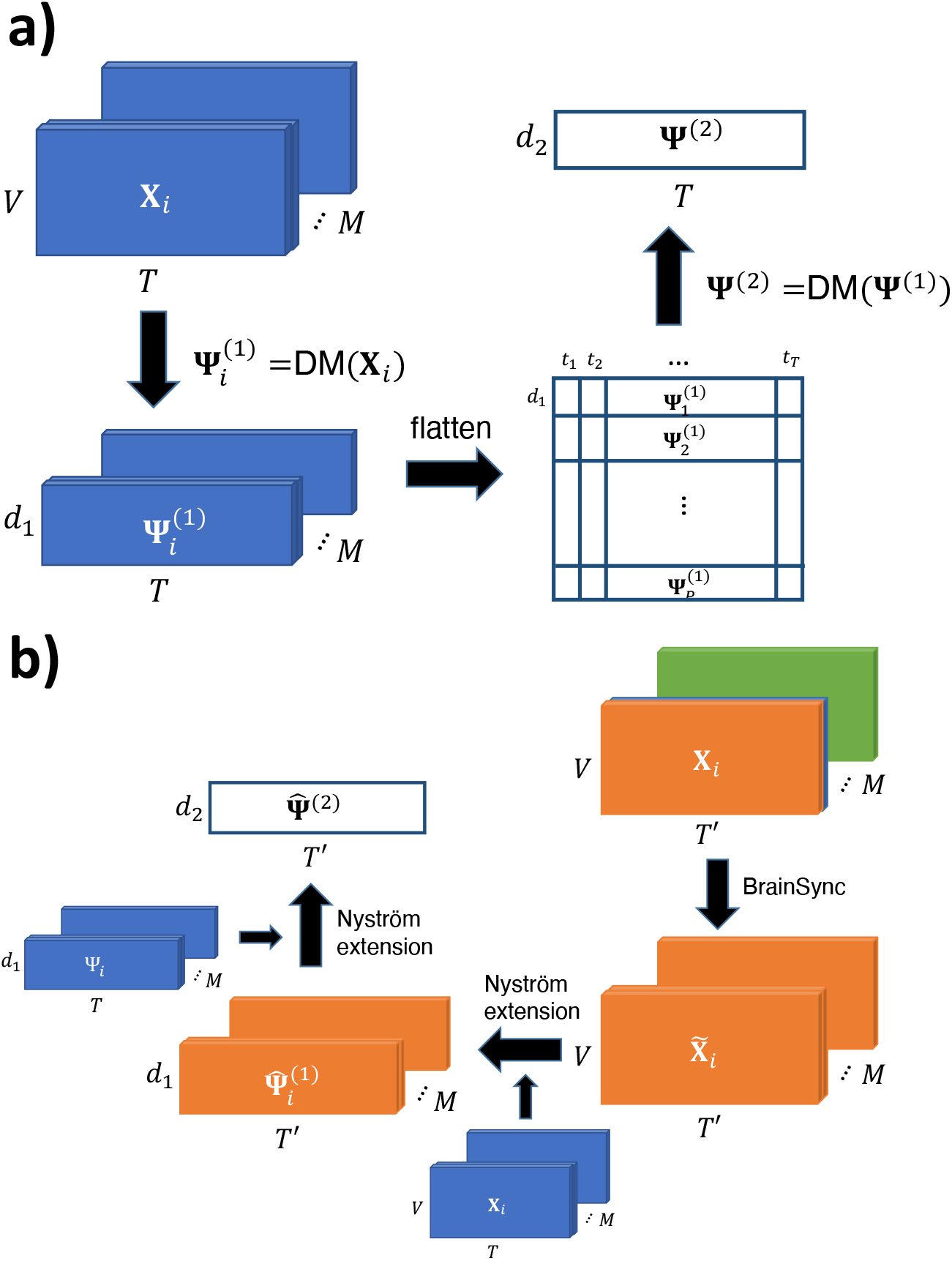
Schematic of manifold learning framework. **a)** 2sDM algorithm framework for time-synchronized multi-individual fMRI time series. **b)** 2-step out-of-sample extension framework with BrainSync for new fMRI time points. Mathematical notations in the figure are the same as those used in the corresponding Methods section.

To reveal the progression of brain dynamics during tasks, we calculated temporal trajectories (Shine et al., 2019) for each task block by connecting points in the embedding in a temporal order. As the tasks involve the same task blocks with repetitions (i.e., WM task consists of interleaved blocks of 0-back and 2-back with the same length), we averaged the time-frames belonging to the same task block to obtain a smoothed representative trajectory of each task. Time frames corresponding to the cue or fixation between tasks blocks were not included.

To summarize the embedding into a more compact and easier to analyze structure, we performed *k*-means clustering based on the first three embedding dimensions to cluster time points sharing similar brain activation patterns into discrete brain states. The Calinski-Harabasz criterion (ratio between the within-cluster dispersion and the between-cluster dispersion) was used to determine the number of clusters, evaluating values of *k* = {2, …, 10} (Caliñski and Harabasz, 1974).

To illustrate that our 2sDM manifold learning framework discovers structure that linear methods cannot recover, we used a 2-step PCA framework, similar to 2sDM. In the first step, a separate PCA is applied to each individual’s fMRI time series X_*i*,,,,_ ∈ ℝ^V×T^, resulting into a *d*_1_-dimensional linear temporal embedding of each individual 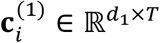, where the first *d*_1_ principal components with the maximum variances are included. Then each individual’s embedding is concatenated along the time dimension form to a matrix 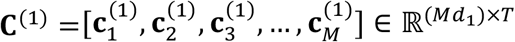. A second-round PCA is performed to further reduce the dimensionality of this concatenated matrix. Each time frame is then embedded into *d*_2_ dimensions and the final time-frame representation with multi-individual similarity is 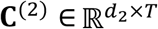. Same as 2sDM, we used *d*_1_ = 7 and *d*_2_ = 3 in our experiment.

### Dynamic connectivity

To relate our task embedding to previously used handcrafted features (Shine et al., 2016), we calculated the participation coefficient *B*_T_ using sliding-window-based functional connectivity and then averaged *B*_T_ across all subjects, as described in previous literature (Shine et al., 2016). In this manuscript, handcrafted features refer to features that are designed manually, such as *B*_T_ that is used here to characterize the integration and segregation pattern of the brain network. The dynamic functional connectivity is calculated by the multiplication of temporal derivatives (MTD; Shine et al., 2015). MTD is calculated as the point-wise product of the temporal derivatives of paired nodes (*i, j*)’s time series:

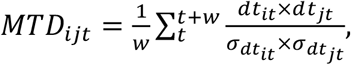

where *d*t_*it*_ = t*s*_*it*_ − t*s*_3*it*−*1*_ is the temporal derivative of node *i* at time t with time series (t*s*), 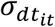 is the standard deviation of the *d*t and *w* is the window length. At each time point, the dynamic functional connectivity is calculated as the averaged MTD over a sliding time window in order to reduce high-frequency noise. We chose the length of the sliding window length *w* to be 15 time points, based on previous literature (Shine et al., 2016).

The participation coefficient *B*_*T*_ characterizes the extent to which a region connects across all modules, where modules are normally defined *a priori* from community detection methods that identify a set of nodes as a module that are more strongly connected to each other than nodes from another set. The participation coefficient for a region *i* at time *T* is calculated as:

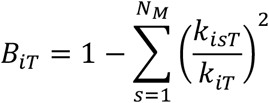

where *k*_*isT*_ is the number of links of node *i* to nodes in module *s* at time *T, k*_*iT*_ is the total degree of node *i* at time *T* and *N*_*M*_ is the number of modules, or canonical networks in our setting. The participation coefficient of a region is therefore close to 1 if its links are uniformly distributed among all the modules and 0 if all its links are within its own module. The whole brain participation coefficient *B*_2_ represents the average of *B*_*iT*_ from each region and thus represents the integration and segregation pattern of the brain. *B*_*T*_ is closer to 1 if our whole brain is more integrated and closer to 0 if our whole brain is more segregated.

### 2-step out-of-sample extension (OOSE) for resting-state fMRI

To investigate the generalization of the task manifold and associated brain states, resting-state data were embedding onto the manifold. One of the challenges in nonlinear dimensionality reduction is to extend new data points to the embedding space. Unlike linear dimensionality reduction methods like PCA, there is no explicit mapping from the original features to the new coordinates. Moreover, appending the new data and redoing the dimensionality reduction is often computationally costly. To deal with this, we specially designed a corresponding 2-step out-of-sample extension (OOSE) framework to embed new time points onto the existing temporal manifold.

The framework is illustrated in Figure 1b. The framework follows a similar two-step structure to 2sDM where the Nyström extension (Fowlkes et al., 2004) (a non-parametric OOSE method, details provided in supplementary materials) is used to approximate the reduced representation of the new time series in each step. Specifically, given new fMRI time series 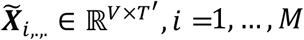 from the same group of individuals used for 2sDM embedding, we first approximate the eigenvectors 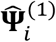 for each individual using Nyström extension. Then we concatenate all the individuals’ eigenvectors 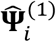 as the new data points and approximate its eigenvectors 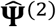 as the final representation.

As the 2sDM algorithm requires the task designs across individuals to be the same, this prevents embedding multi-individual resting-state fMRI timeseries directly, which is also a problem for any other scans that are not time-synchronized, e.g., the language task in the HCP dataset. To synchronize different individual’s time series, we used BrainSync, a framework that synchronizes fMRI time series across individuals (Joshi et al., 2018). BrainSync synchronizes one individual’s time series data ***Y*** ∈ ℝ^V×*T*^ to another reference individual **X**∈ ℝ^V×*T*^ by finding an optimal orthogonal transformation that minimizes summed moment-to-moment squared error O^*s*^ = arg m*i*n_O∈O(*T)*_‖X−*Y*O^*t*^‖^2^. The problem can be solved by the Kabsch algorithm (Kabsch, 1976). The *T* × *T* cross-correlation matrix **X**^*t*^*Y* is first formed and its singular value decomposition can be calculated as **X**^*t*^***Y*** = **UΣV**^*t*,^. The optimal O^@^ can be found by O^*s*^ = **UV**^*t*^ and ***Y*** can be synchronized to **X** by 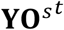. By choosing a random individual as the reference, BrainSync was applied to all the other individuals and their time series were synchronized to the reference individual. After synchronizing across individuals, we then used the 2-step OOSE framework to extend them onto the task manifold and find the temporal representation of resting-state fMRI for the reference individual.

To validate the 2-step OOSE framework, we used the task fMRI data to cross-validate the accuracy of the OOSE framework. Using leave-one-task-out cross-validation, a single task was held out when generating the 2sDM manifold. The left-out task was then embedded in the new manifold using our OOSE framework and compared with the original embedding created using all tasks. If the held-out task’s extended coordinates are similar to the coordinates of the original embedding, it suggests that the OOSE framework is accurate.

### Characterizing changes in brain states

By utilizing the temporal order of time points, we characterized the brain dynamics across the four brain states by state transition probability and dwell time. State transition probabilities were calculated based on the temporally adjacent time points’ brain states. From these state transition probabilities, a stochastic matrix and the dwelling times (i.e., the stationary probability distribution of the stochastic matrix) were calculated and visualized as Markov chain models.

The stationary distribution of the Markov transition matrix **P**_**trans**_ is defined as the distribution that does not change under application of the transition matrix **π** = **πP**_**trans**_, which is the left eigenvector of **P**_**trans**_. It represents the distribution to which the Markov process converges. It was used in our experiment to represent the dwell-time distribution of discrete brain states. As tasks putatively put a participant into certain states (as opposed to the unconstrained nature of the resting state), we investigated differences in the temporal dynamics of state switching during task and rest. We calculated entropy—a measure of the randomness—of the transition probability from one brain state to the other states. Entropy of a discrete probability distribution measures the uncertainty of the outcome. It is calculated as the negative expectation of the logarithm of the probability mass function’s value *S* = −Σ_*i*_*P*_*i*_ log *P*_*i*_ = −*E*_*p*_[log *P*]. In our experiment, entropy of the brain state transition probability was used to assess the randomness of brain state transitioning with lower entropy representing more easy-to-predict brain state transition dynamics. Greater entropy indicates a less predictable transition from one state to another.

### Experimental Design and Statistical Analysis

No statistical methods were used to predetermine sample sizes. Other than the stated exclusion criteria for motion and complete imaging data, no participants and data points were excluded from the analysis. Following exclusion for motion, there was no significant correlation between motion and the embedding dimension. Parametric statistics (e.g., *t*-test, correlation, and chi-squared test) were used when appropriate.

### Data availability

The HCP data used in this study are publicly available from the ConnectomeDB database (https://db.humanconnectome.org). The CNP data used in this study are publicly available from OpenNeuro.org (https://openneuro.org/datasets/ds000030). MATLAB scripts to run the 2sDM analyses can be found at (https://github.com/carricky/2sDM). BioImage Suite tools used for analysis can be accessed at (https://bioimagesuiteweb.github.io/).

## Results

### Brain dynamics during tasks embed onto a low-dimensional space

Although each task is different in many ways, individual time points in the fMRI data from all tasks mapped onto a single low-dimensional manifold (Figure 2a). Compared with the common goal of other low-dimensional embedding results, the advantage of our results is not in separating different task scans apart. Instead, we find a global representation across multiple tasks that positioned tasks with similar cognitive loads together. By embedding multiple tasks together, rather than in isolation, the closeness of different blocks and tasks in the manifold suggest that similar, recurring patterns of brain dynamics exist across a variety of tasks. For example, in the manifold, the 2-back blocks of the WM task are significantly (t = 201,9, *p* < 0,01, d, f, = 175,102) closer to time points from the gambling task (Euclidean distance: 0,0258 ± 0,0096) than the 0-back blocks of the WM task (Euclidean distance: 0,0355 ± 0,0100), despite the fact that the 2-back and 0-back blocks were collected in the same fMRI run. The 2-back blocks of the WM task and the gambling task both entail a higher cognitive load. In contrast, the 0-back blocks of WM task overlap with the motor task. These tasks are simpler response tasks and less cognitively demanding. Overall, these time points are positioned based on the similarity of the cognitive load at that time point, instead of by task.

**Figure 2.**
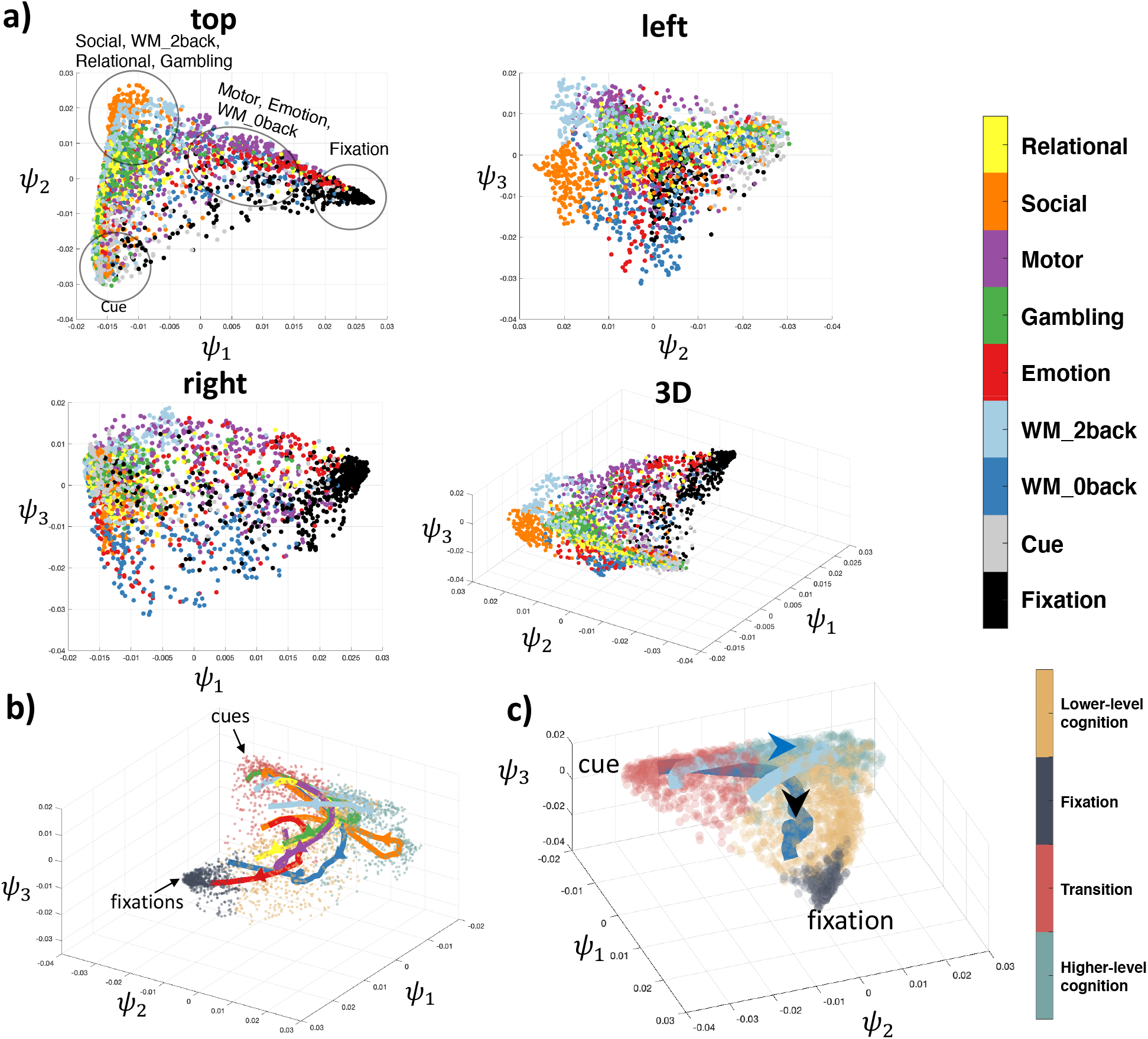
Nonlinear embedding of fMRI time series data. **a)** 2sDM embedding of 6 tasks (relational, social, motor, gambling, emotion, working memory 2-back, and working memory 0-back) from the HCP dataset. Four different views of the manifold are shown. Each point in these subplots represents a single time point and is colored by the task type. **b)** Averaged temporal trajectory of each task with the embedding colored by the corresponding brain state as the background. **c)** WM task’s 0-back and 2-back task blocks visualized separately with major cues and fixations points annotated. Arrows show the progression direction of the trajectory. Trajectory in b) and c) uses the same colormap as a).

For all tasks, the average trajectories from each task are found to start near the corner where cues (task cues preceding each task block) reside and end in the other corner where fixation blocks reside. These smooth trajectories indicate that the embedding preserves proper temporal associations between blocks when arranging time points in discrete states. As can be expected, the paths of these temporal trajectories depend on the cognitive load of the task block. For example, the 2-back task traverses through the upper part of the manifold (higher value in terms of ψ_3_), and, in contrast, the 0-back task traverses through the lower part of the manifold (Figure 2b). Moreover, as can be seen from the top 20 eigenvalues of the diffusion matrix the spectrum decays rapidly, which suggests that the data is low-dimensional (Figure S1).

When projecting task fMRI time-frames into 3D space using the first three coordinates of PCA, no clear structure is shown from the embedding (Figure 3). The fact that 2sDM discovered the manifold structure, while PCA could not, validates the usage of nonlinear manifold learning (more detailed comparison between 2-step PCA and 2sDM embeddings are included in the supplementary materials, Figure S4-S7).

**Figure 3.**
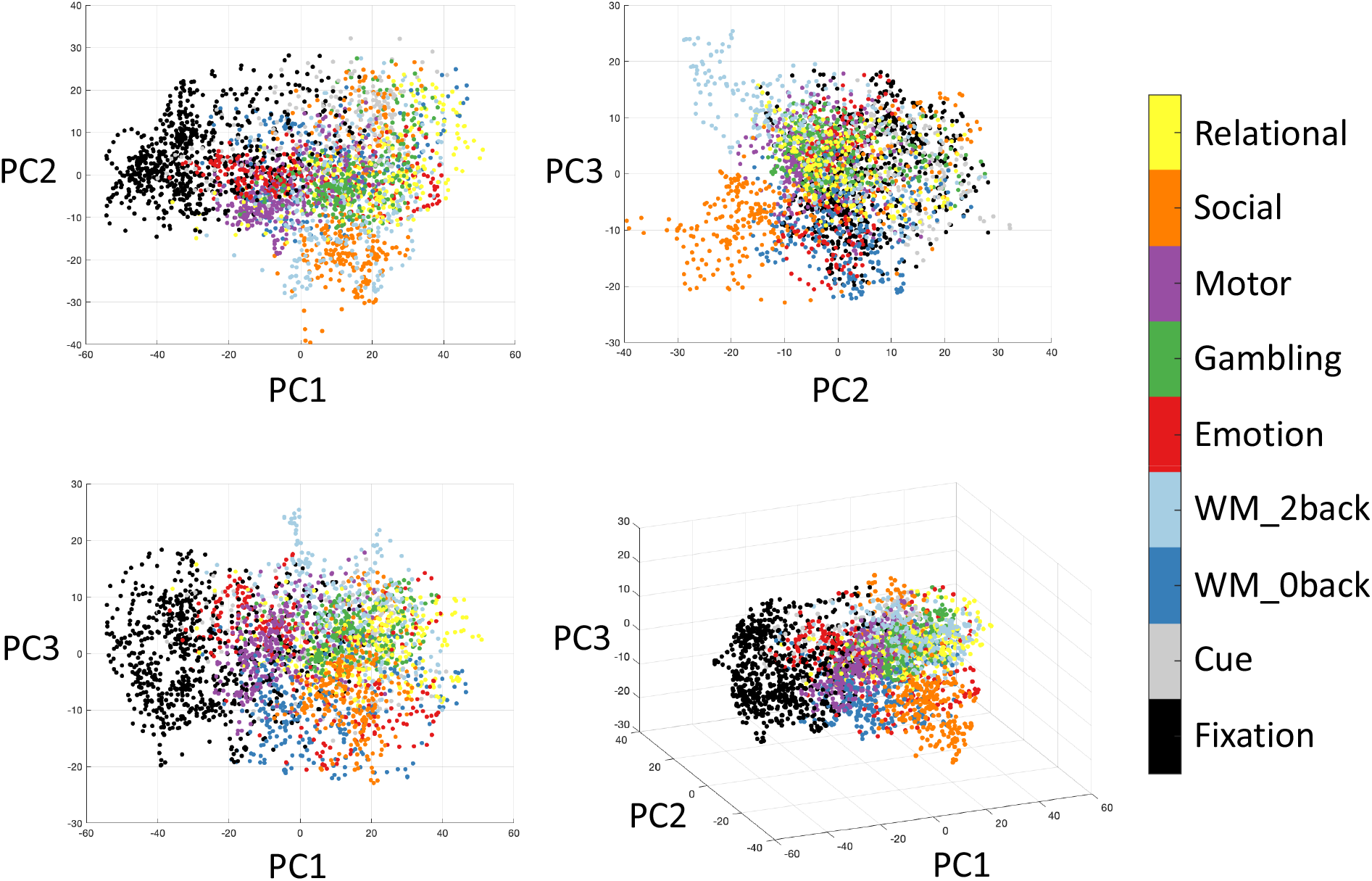
2-step PCA embedding from the HCP dataset. Unlike the nonlinear embeddings, shown in Figure 2, no clear structure is seen for the linear embedding, which validates the usage of nonlinear manifold learning.

### Task embedding captures handcrafted features in an unsupervised manner

In Figure 4a, each time point in our task embedding is colored by its subject-averaged *B*_*T*_, showing a clear pattern of decreasing *B*_*T*_ starting from the top left corner of the embedding; higher *B*_*T*_ at the top of the embedding (i.e., high cognitive load tasks such as social, 2back, relational and gambling) indicates time points of higher integration and lower *B*_*T*_ at the tails of the embedding (i.e., cues and fixations) indicates time points of higher segregation (*r*(*z, B*_*T*_) = 0,610, d, f, = 3018, *p* < 0,01, where *z* is the projection coordinates of points onto the diagonal of the triangular embedding; Figure 4b).

**Figure 4.**
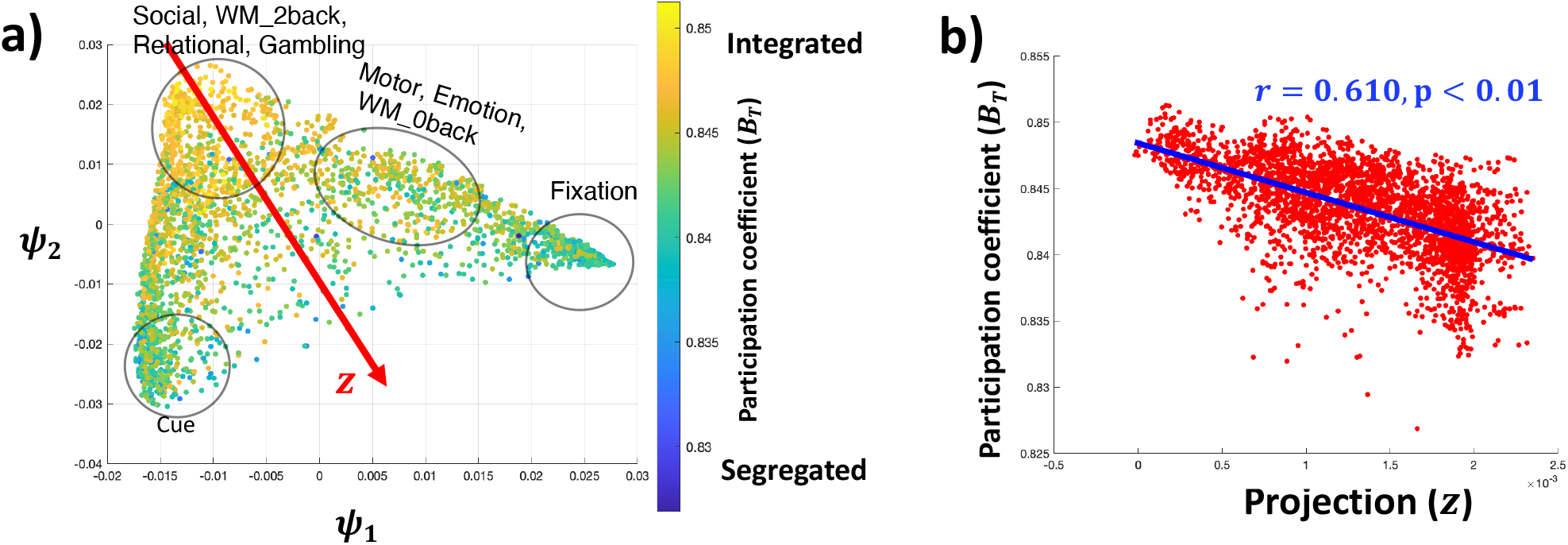
2sDM embedding is related with global integration and segregation. **a)** 2sDM embedding in HCP dataset colored by the time-resolved *B*_T_. **b)** Scatter plot of the *B*_T_ with the projection onto the diagonal of the embedding structure (z). Correlation of z with *B*_T_ is shown with a line of best fit. Projection direction z was determined manually as the approximate diagonal direction of the embedding.

### Operationalizing discrete, recurring brain states from task dynamics

When clustering the task embedding, *k* = 4 gives the largest Calinski-Harabasz score among a range, suggesting that the embedding has a clear interpretable structure (Figure S6). Based on the task contents of the temporal clusters, we labeled the four brain states as: fixation, transition, lower-level cognition, and higher-level cognition. Functionally reasonable and distinct patterns of activation during the different states are observed, e.g., canonical patterns of default mode network activity for the fixation state (Figure 5a). To relate these brain states to previous handcrafted features, we calculated the average *B*_T_ for each brain state (Figure 5b). The four states followed the expected patterns of integration and segregation, with the higher-level cognition state showing the greatest integration (*t* = 3,01, *p* < 0,01, d, f, = 1596) and the fixation state showing the greatest segregation (*t* = 2,39, *p* < 0,01, d, f, = 1420). The clustering results are similar with an increased number of clusters or of embedding dimensions.

**Figure 5.**
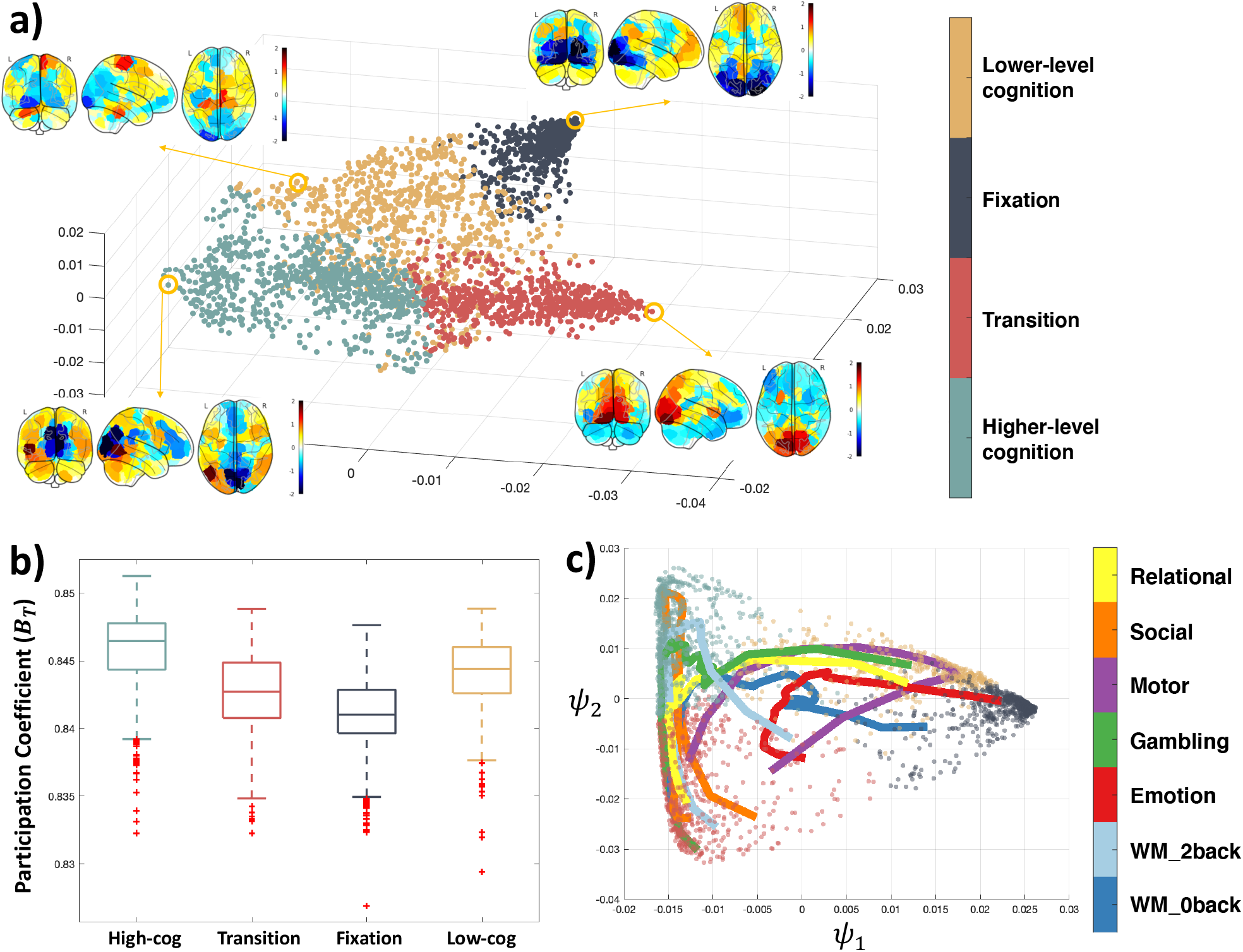
Brain states during tasks. **a)** Results of *k*-means clustering of the task manifold. Averaged brain activation patterns across subjects in the circled representative time points are shown for each brain state. **b)** *B*_T_ averaged over all the time points in each brain state. **c)** Twodimensional view of task trajectories with the embedding points. Trajectories are colored by each task and data points are colored by the brain states as in a).

With the help of the four brain states, the dynamic trajectories can further reveal each task’s cognitive process (Figure 5c). For example, the motor task’s trajectory reveals a dynamic cognitive process as following: in the beginning, the individuals start from the cue state which is the common starting state across the other tasks. Then the individuals briefly enter the high-cog state, but not deep in the state and finally enter and stay in the low-cog state. Actually, it also reveals that on average, individuals wander towards the fixation state in the middle of the task block, suggesting a fatigue or practice effect. And towards the end of the task block, individuals return deep into the low-cog state and move towards the cue state for the next task block to start.

Even for tasks like relational and social tasks that both require a certain level of high-level cognitive ability (Shine et al., 2016), there are differences that can be revealed by the trajectories (Figure 5c). The relational task starts from the transition cluster, then entered the higher-level cognition cluster and ends in the low-cog state, which suggests a lack of high-level cognitive ability involvement (adaptive to the task design) in the later stage of the relational task blocks. In comparison, the social task starts near the transition cluster, goes deep into the high-cog state and returns to the transition state near the end of the task which suggests a constant requirement of higher-level cognitive ability. This trajectory view of each task enables a better understanding of the cognitive process and can also help in the future task designs.

The transitions between states are similar for all tasks except for the motor task (which had a high probability of transiting into the lower-level cognition state and out of the higher-level cognition state; Figure 6a). Except for the WM task, which contains an equal proportion of high (2-back) and low (0-back) cognitive loads), dwell times for the four states exhibit a non-uniform distribution (*χ*^2^ > 16,3, d, f, = 3, *p* < 0,001; Figure 6b), indicating participants spent most of their time in certain limited states in a task-specific manner. For example, the lower-level cognition state occurs most frequently in the motor task, while the higher-level cognitive state dominates in social task time points.

**Figure 6.**
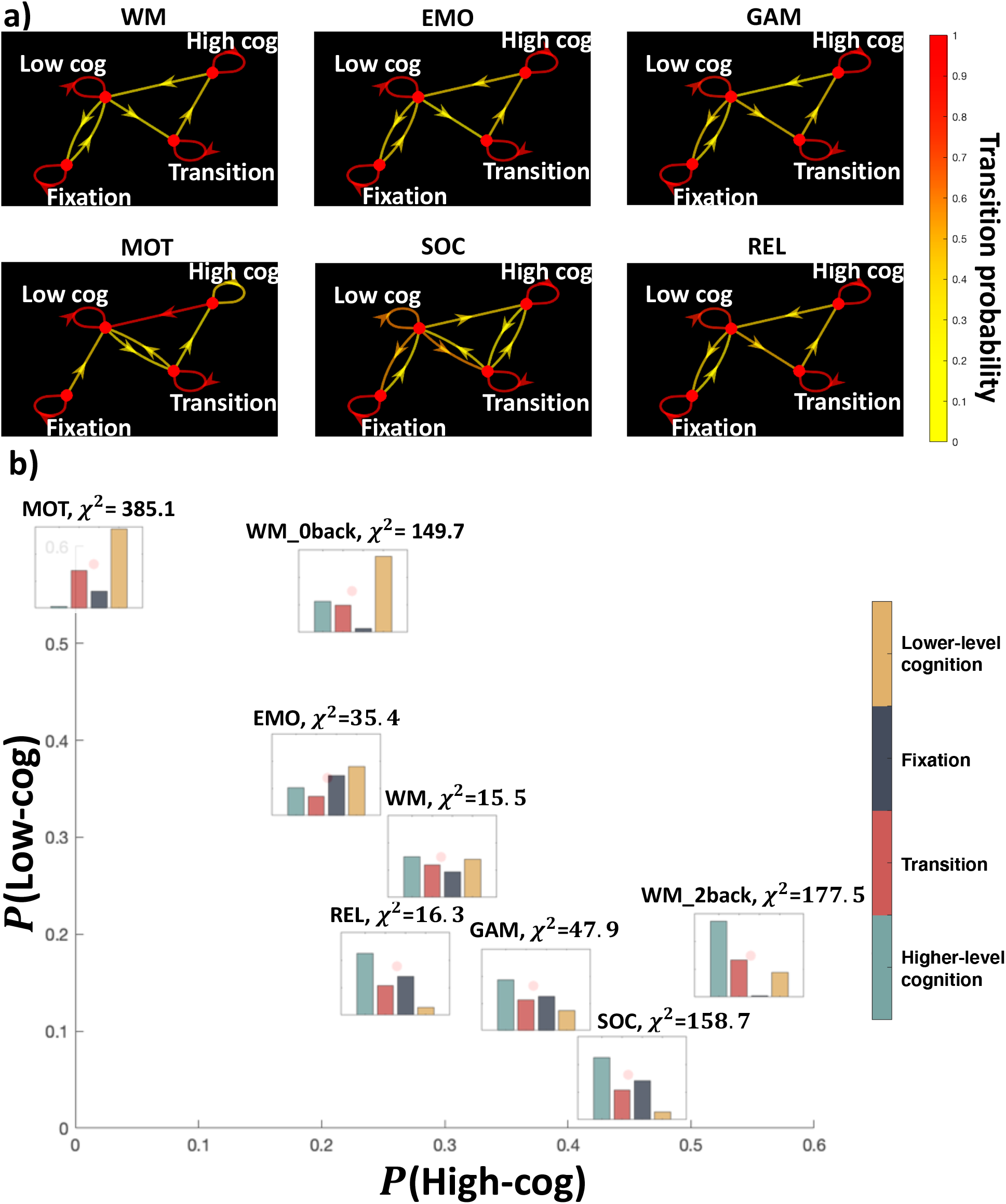
Brain state dynamics differ between tasks. **a)** Brain state dynamics visualized as the Markov chain. Transition probability is visualized by the color of the directed edges. **b)** Stationary distribution probability visualized for each task and positioned by the proportion of higher-level cognition and lower-level cognition brain states. Chi-square test result against the uniform distribution is also shown.

### Brain dynamics during rest embed onto the same recurring brain states which appeared during tasks

Once embedded onto the task manifold, time points from the resting-state data spread across the whole manifold, including parts of the manifold corresponding to higher cognitive loads (Figure 7a). To quantify the distribution of states during rest, we assigned each resting-state time point to one of the four previously identified brain states based on the brain state of the nearest task time point. As with the task data, we next calculated the brain state dwell time distribution across the entire resting-state scan (Figure 7b). A non-uniform dwell-time distribution is discovered, with fixation and transition states having a higher proportion of time points than the cognitive states (*χ*^2^ = 205, d, f, = 3, *p* < 0,001). Except for the lower-level cognition and the transition states in the social task (which have very few time points to robustly calculate entropy, see Figure 7c), all states exhibit higher entropy in the resting state than during a given task.

**Figure 7.**
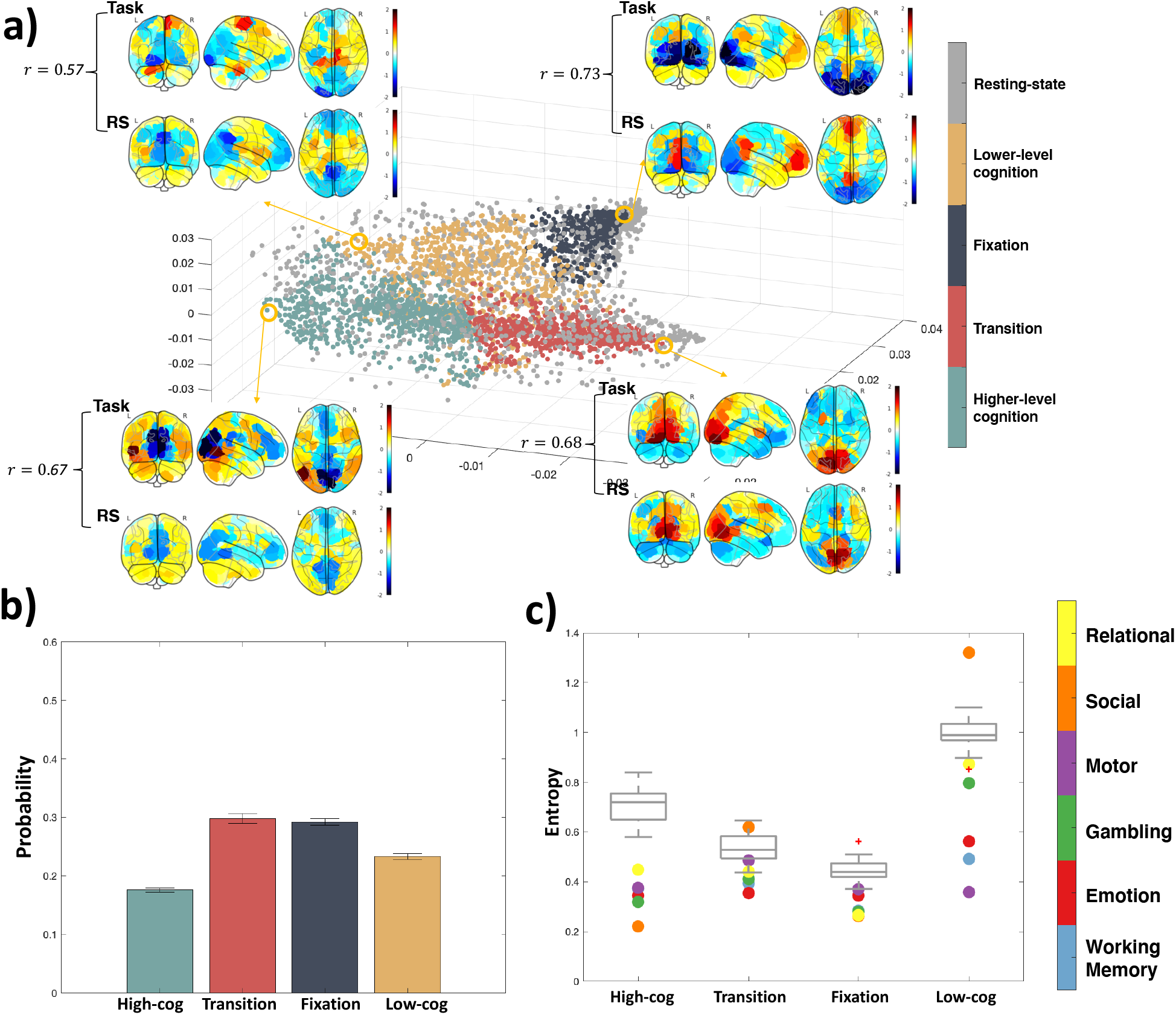
Resting-state extended onto the task manifold. **a)** Representative task activation patterns of each state and the neighboring resting-state activation pattern are visualized. Correlation of the activation between task and rest is calculated with higher correlation representing more accurate out-of-sample extension. **b)** Stationary probability distribution of the four brain states during resting state. **c)** Entropy of each brain state’s transition probability in different tasks. Dots are colored by tasks they represent, and the grey box plot shows the entropy values of resting state with BrainSync (see Methods) referenced to different individuals.

In Figure S2, we plot the extension of the WM task. The 2-back and 0-back task blocks go to the correct higher-level cognition or lower-level cognition state respectively, while the fixation and cue time frames are also located in the correct brain states. The correlation between the extended coordinates and the coordinates from the original embedding was highly significant (*r* = 0,939, *p* < 0,001). Holding out the other tasks produced similar results as the WM task.

### Replication of embedding

Notably, we replicated the dimensionality reduction result using participants from the CNP dataset. A similar low-dimensional structure, brain states, and association with *B*_*T*_ (*r*(*ψ*_2_, *B*_*T*_) = 0,30, *p* < 0,01, *df* = 1007) were found, verifying the robustness of the observed embeddings (Figure 8). Moreover, the same task scans from the schizophrenia cohorts were also embedded separately and found to be similar to the embedding from the HCP dataset and healthy control cohorts in the CNP dataset (Figure S8). This laid foundation for the downstream brain dynamics analysis (resting-state brain dynamics) that would be based on brain states as similar brain states could be identified in both groups.

**Figure 8.**
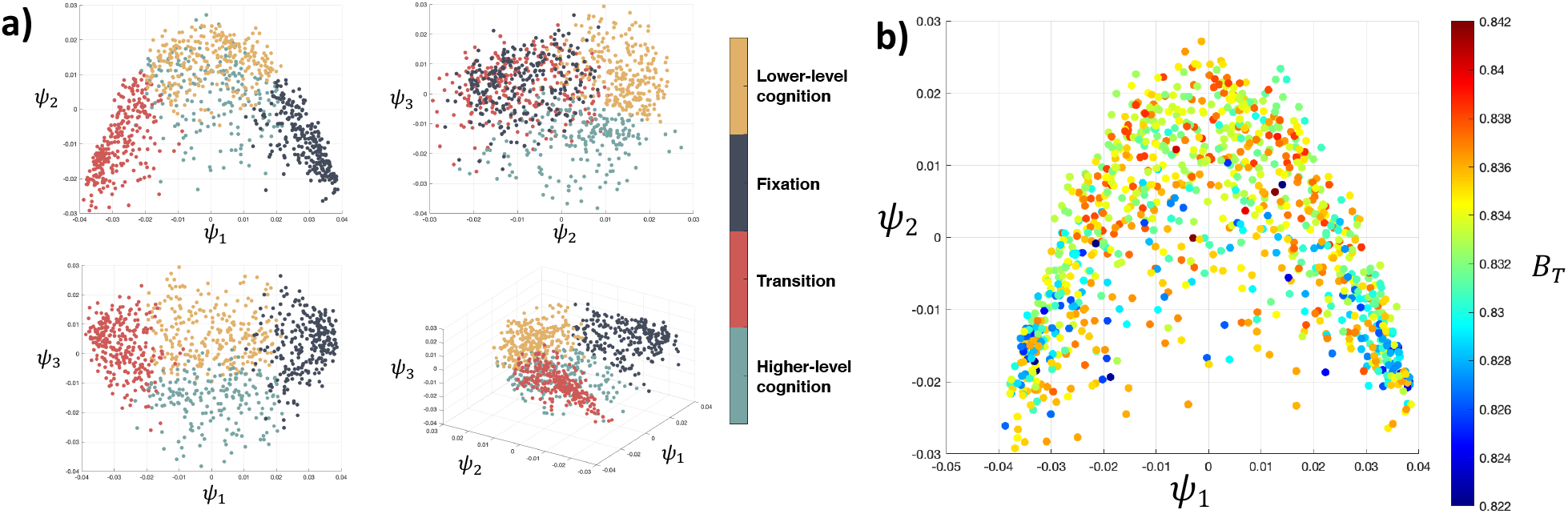
**a)** 2sDM embedding and *k*-means clustering result of CNP dataset. **b)** Embedding with the first 2 dimensions of 2sDM in CNP dataset, colored by the corresponding *B*_T_ with the same colormap.

## Discussion

Using a novel manifold learning framework, we demonstrate that fMRI data from different tasks span the same low-dimensional embedding (i.e., brain states). In other words, moment-to-moment dynamics from any of these tasks group into the same small number of representative patterns that are hidden from direct observation. To recover this embedding, we employed nonlinear methods (e.g., 2-step Diffusion Maps—2sDM) to project the fMRI data onto a manifold than would be possible using linear methods only. The embedding maintained proper temporal progression of the tasks, revealing brain states and temporal dynamics of changes in network integration. Further, we demonstrate that resting-state data project onto the same task embedding using a specially designed out-of-sample-extension method, indicating similar brain states are present. Finally, we validate this embedding using an independent dataset.

Several other publications have organized the temporal dynamics of the brain into a low dimension space or into distinct brain states (Allen et al., 2014b; Saggar et al., 2018; Vidaurre et al., 2017) using data from resting-state or a single task to construct the embedding (Gallego et al., 2017; Shine et al., 2019). Together, these works suggest that a low-dimensional structure exists; however, it is unclear how these structures adapt to diverse cognitive loads. By projecting a rich repertoire of task data into a single manifold, we show that, across different tasks, parts of the embedding (i.e. brain states) are well characterized by network segregation (i.e. communication mainly within brain networks) and integration (i.e. communication mainly across diverse brain networks) (Deco et al., 2015). Overall, the discrete states and association with network segregation/integration suggest that our embedding finds an intrinsic, latent structure of brain dynamics.

These results are in line with the theory that the brain is able to reconfigure its large-scale organization dynamically either between different cognitive tasks or within resting-state (Cohen and D’Esposito, 2016; Shine et al., 2016). Further, they emphasize that this reconfiguration is shared across different cognitive loads and, importantly, resting-state. In other words, the same highly integrated state that characterizes a cognitively demanding task, such as a 2-back WM task, can be observed during resting-states and less cognitively demanding tasks, just with less frequency. These states can also be viewed from a dynamic system perspective (Taghia et al., 2018). As clustering based on the eigenvectors of the normalized graph Laplacian has been used to find meta-stable state in the stochastic dynamical systems (Huisinga et al., 1999), the four brain states defined from the task scan can also be viewed as four different metastable states. Further, the temporal trajectories can separate different portions of tasks based on cognitive demand, suggesting a potential utility of the embedding for other downstream analyses of brain dynamics.

In line with this, the dynamics between states, rather than within brain states themselves, appear to be the key distinguishing factor between task and rest. In support of this, how the brain transitions between different states is dependent on the task being performed and is less predictable in resting-state compared to tasks. Executing a task limits the transitions between states; while, during resting-state, the brain can more liberally traverse through different states. Though speculative, these results offer an explanation as to why task connectivity data is better at identifying individuals and subsequent predicting behaviors than resting-state connectivity data (Finn et al., 2017; Greene et al., 2018). Together, while the resting state may exhibit similar states as observed during task, the temporal dynamics of switching states are less predictable in resting state compared to task.

Previous work demonstrates that brain networks fluctuate between states of low and high global integration during tasks as characterized by the participation coefficient (*B*_*T*_) from sliding-window functional connectivity. Tasks requiring higher cognitive loads, such as the 2-back condition in the WM task, exhibit greater integration while less cognitive load, such as the motor task, exhibits lower integration (Shine et al., 2016). A key drawback of these results is that they rely on two intermediate steps (e.g., the method used to construct dynamic functional connectivity and topological metrics to study), rather than the learned features from unsupervised methods. Together, our results suggest that the task embedding reveals latent information about changes in network topology without the need for handcrafted features. For example, each task can be effectively characterized from the proportion of time spent in lower-level and higher-level cognition states creating a similar ordering of task (see Figure 6b) as in (Shine et al., 2016).

While resting-state fMRI is a powerful tool to map the functional organization of the brain, inherent limitations exist. Resting-state is often conceptualized as a single task state. Though emerging data, including our results, suggest that resting-state is not one single, monolithic state, but rather a collection of multiple states associated with different cognitive loads that also appear during tasks (Vidaurre et al., 2017). For example, while the majority of resting-state time points cluster into a single part of the manifold (such as the fixation blocks, which putatively are the most like “rest”), nearly a third of the time points more closely match cognitive states. Perhaps, more importantly different groups may have differences in “performing” rest (Buckner et al., 2013). How best to interpret changes in resting-state connectivity in the presence of group differences in dynamics is still an open question.

A key strength of our embedding framework is its data-driven nature. Although the only inputs are time-courses from task fMRI data, we demonstrated that the embedding coordinates can reveal topological information originally found using dynamic functional connectivity methods (Shine et al., 2016). This brain topology was found without specifying common modeling choices in dynamic functional connectivity or fMRI, in general, such as how to model the functional connectivity (i.e., statistical interdependence of signals) between brain regions, an underlying graph/network, or even information about task stimuli (e.g., block lengths). As a multitude of methodological choices have been proposed to analyses (Calhoun et al., 2014; Hutchison et al., 2013a) (e.g., ways of estimating connectivity (Allen et al., 2014a; Chang and Glover, 2010; Shine et al., 2015), constructing a weighted or unweighted graph (Rubinov and Sporns, 2010), specific graph theory measures (Honey et al., 2007; Meunier et al., 2010; Shine et al., 2016; Sizemore and Bassett, 2018), our embedding framework provides an end-to-end, data-driven approach without the need for modeling choices to investigate brain dynamics. More generally, handcrafted features are being substituted by more automatic feature learning-based nonlinear methods such as deep learning and nonlinear embedding methods (Hamilton et al., 2017). Our results show a specific scenario in which “let the data speak for itself” is an achievable option for modeling fMRI data.

A limitation of this work is that the embedding can only “look under the light.” That is to say that, while a rich amount of task data was needed to create the embedding, we could not include every possible task in creating the embedding. Indeed, it is highly likely that many more than four brain states exist and that we haven’t detected every single one. A finer grade delineation of states, probably through further advancement in non-linear embedding methods, is a needed future direction of work. Moreover, although here brain states are defined based on the *k*-means clustering result, it does not rule out other ways to define brain states. For example, at each time point, the brain can also be modeled as being at different states with distinct probabilities (Vidaurre et al., 2017), which can be achieved by a fuzzy-clustering algorithm. Moreover, the brain state can also be characterized by the temporal trajectory where trajectory clustering technique can be used to cluster trajectories into trajectory-based brain states, which takes account the temporal information of the embedding (Lee et al., 2007). The *k*-means clustering way of defining brain state is only one of the ways to summarize information of the embedding and serves as a proof-of-concept that our embedding contains information that is relevant to brain dynamics. Nevertheless, the observed task embedding was similar across two different input datasets with different tasks, suggesting that embedding is general to factors such as scanner, task, processing, and sample size.

One of the assumptions of 2sDM is that the time frames from all individuals are temporally aligned so that a group-average embedding of the time frames can be obtained. However, this does not rule out the applicability of the task scans that has different task block lengths/orders across individuals (e.g., language task in the HCP dataset) or the resting-state scans, which we have demonstrated in the paper by applying BrainSync. Thus, task scans with distinct block lengths/orders can also be embedded with 2sDM by applying BrainSync first. It is worth noting that as BrainSync requires a specific individual chosen as the reference, by aligning all the other individuals to the same selected individual, the group-average embedding then will approximate a cleaner temporal embedding of the selected individual, which can be used to investigate individual-level dynamics.

The ability to use data-driven methods to clearly identify a low-dimensional space of brain dynamics, regardless of how the brain is engaged during imaging, indicates that these brain dynamics are robust and reliable across conditions in addition to being unique. Together, these advances suggest that analysis of individual fMRI data from multiple cognitive tasks in a low-dimensional space is possible, and indeed, desirable.

## Acknowledgments

Data were provided in part by the Human Connectome Project, WU-Minn Consortium (Principal Investigators: David Van Essen and Kamil Ugurbil; 1U54MH091657 funded by the 16 NIH Institutes and Centers that support the NIH Blueprint for Neuroscience Research; and by the McDonnell Center for Systems Neuroscience at Washington University) and the Consortium for Neuropsychiatric Phenomics (NIH Roadmap for Medical Research grants UL1-DE019580, RL1MH083268, RL1MH083269, RL1DA024853, RL1MH083270, RL1LM009833, PL1MH083271, and PL1NS062410).

