## Supplementary material for "Non-linear manifold learning in fMRI uncovers a low-dimensional space of brain dynamics"

### Spectrum of the data

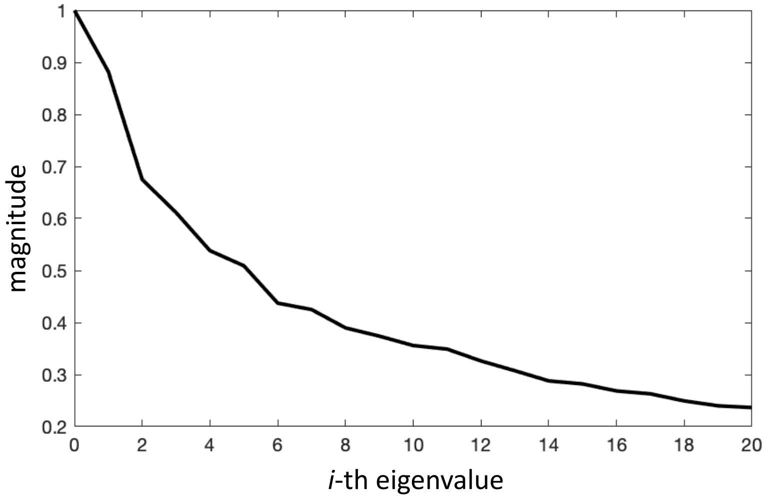


**Figure S1. The top 20 eigenvalues of the diffusion matrix for the fMRI data.** The spectrum decays rapidly, suggesting that the data is low-dimensional (Coifman and Lafon, 2006).

### Validity of out-of-sample extension

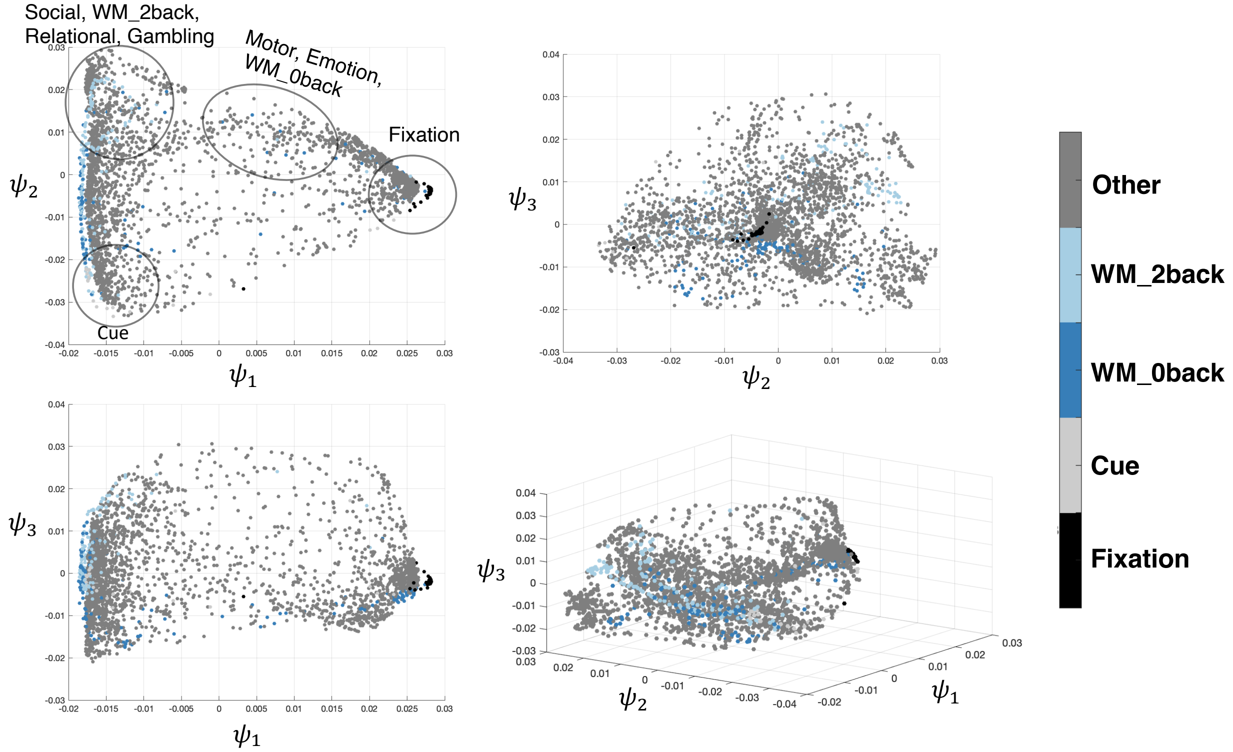


**Figure S2. Extension of WM task's LR session.** The 2-back and 0-back task blocks go to the correct higher-level cognition or lower-level cognition state respectively, while the fixation and cue time frames are also located in the correct brain states. The correlation between the extended coordinates and the coordinates from the original embedding was highly significant, $r = 0.939, p < 0.001$.

### Choice of d1 and d2

The parameter $d_1$ determines the number of dimensions to keep for the first-step subject-wise embeddings while $d_2$ determines the total number of dimensions to keep in the final embedding. As each dimension of the diffusion embedding is calculated based on the eigenvectors of the random walk matrix, adding more coordinates (increase $d_1$ or $d_2$) won't affect the previous coordinates.

In the paper, $d_1$ is set as 7, so that the first 7 dimensions of the first-step embeddings are used to calculate the group-wise affinity matrix. To test $d_1$'s influence, we further test $d_1$ as [3, 5, 9, 11] and correlate the final embedding with the embedding we use in the paper, which is generated with $d_1 = 7$.

In the figure below (Figure S3), we show the correlation of each coordinate generated with $d_1 = 7$ compared with the coordinates generated with other $d_1$ parameters. The first 7 coordinates of the final embedding are tested. As shown in the figure, the first 2 coordinates from all the embeddings are also identical ($r > 0.98$) regardless of different $d_1$ values, suggesting the robustness of the patterns revealed with the first 2 coordinates.

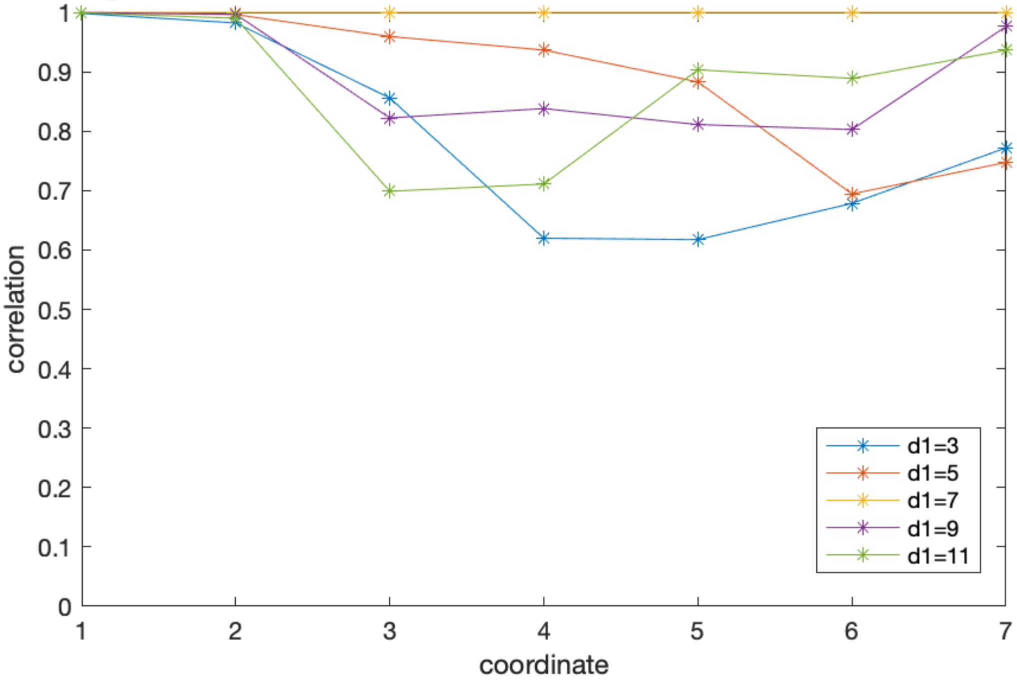


**Figure S3 Influence of different $d_1$ parameter on the final 2sDM embedding results.** Correlation of each coordinate generated with $d_1 = 7$ compared with the coordinates generated with other d1 parameters is shown.

Although the coordinate difference is bigger after the 3rd coordinate, it is expected as when more dimensions are included in the single subject embedding, the more noises and local information are also included. Overall, those coordinates are still highly correlated ($r > 0.6$) with the coordinates we use in the paper that are generated with $d_1 = 7$.

As the first 7 dimensions in the final embedding are shown to be relatively robust, we don't use all of them as the first 3 dimensions ($d_2 = 3$) have provided us interesting results to illustrate the power of the framework and are easier to visualize. From the spectrum of the diffusion matrix (Figure S1), the first 2 dimensions have dominant influence.

As they are also shown to be robust under different $d_1$ parameters, our conclusions in the paper are not impacted by the hyperparameter choices.

### Comparison to 2-step PCA

Although our 2sDM framework is not fixed to only the diffusion maps algorithm (Monti et al., 2017), we compare the 2sDM results with its linear comparison, 2-step PCA (2sPCA), in more details here.

*Low-dimensional embedding*

Unlike the related low-dimensional embedding works that only involved two or three tasks (Monti et al., 2017; Venkatesh et al., 2019), in this work, as we try to embed multiple different tasks (6 tasks with sub-block tasks in the HCP dataset) together, our goal is no longer separating different tasks. Instead, we aim to cluster different tasks together in a meaningful way to reveal the common factors that drive different tasks. Therefore, when we compare the embedding from 2sPCA vs 2sDM, we examine whether 2sPCA also reveals a meaningful embedding structure. However, as shown in Figure S4, without the colors as prior, it is harder to infer the structure from the 2sPCA embedding. Although different tasks are separated (Figure 3), the cue and fixation time points don't form two separate corners as in the 2sDM embedding, which could make the downstream analysis less interpretable (illustrated in the following trajectory analysis). Moreover, it is obvious to analyze the three corners of the 2sDM embedding at first as they are the anchors of the embeddings while harder to choose similar points in the 2sPCA embedding.

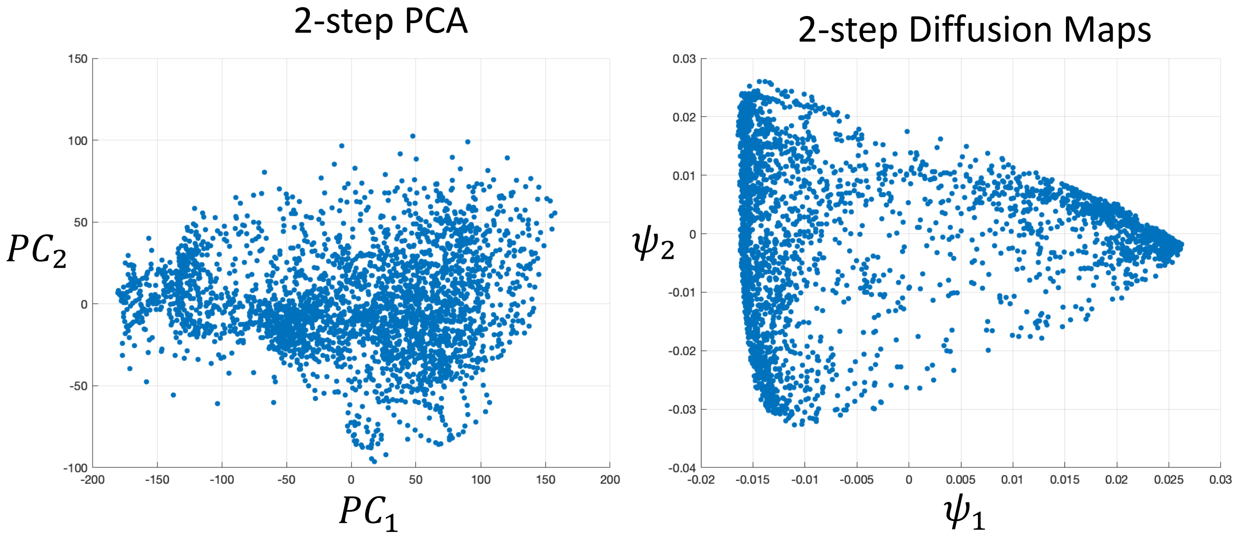


**Figure S4** Comparison of 2-step PCA embedding with 2-step Diffusion Maps in the first 2 embedding coordinates without the points colored by task block types.

*Temporal trajectory analysis*

As we illustrate in the main paper that the low-dimensional temporal trajectories formed with each task conditions reveal the dynamic cognitive processes, the 2sPCA trajectories are less informative. In the Figure S5, we show the average trajectory for each task. As the 2sDM embedding clusters the cues and fixations in separate corners, the 2sDM trajectories are easier to interpret and compare across tasks. In comparison, as the fixation and cues are more scattered in the 2-step PCA embedding, the cross-task comparison is more difficult. For example, we can infer that for the Motor task, the 2sDM trajectory reveals the cognitive process where in the beginning, the individuals start from the cue state which is the common starting state across the other tasks. Then the individuals briefly enter the high-cognitive state, but not deep in the state and finally enter and stay in the low-cognitive state. It also reveals that on average, individuals wander towards the fixation state in the middle of the task block, suggesting a fatigue or practice effect. And towards the end of the task block, individuals return deep into the low-cognitive state and move towards the cue state for the next task block to start. However, none of these analyses will be obvious for the 2-step PCA trajectory as the motor task trajectory starts in a different location and has a different progression pattern from other tasks.

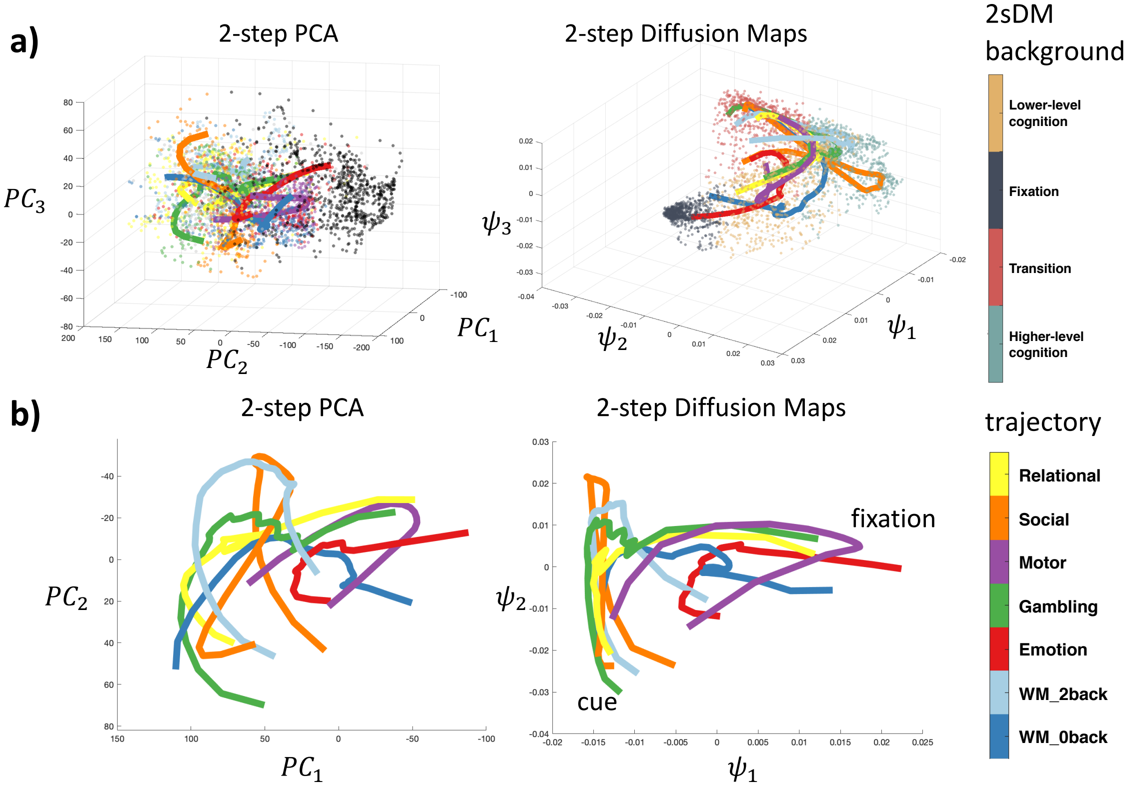


**Figure S5 Trajectory comparison of 2sPCA and 2sDM. a)** 3-dimensional view with the embedded time points in background. 2-step PCA's background time points are colored by the task type (same colormap as the trajectory) and 2-step Diffusion Maps' background time points are colored by the brain states. **b)** 2-dimensional view of the trajectories only. The first two dimensions of the embedding are used. Figures of 2-dimensional views are rotated to match with each other.

*Brain state clustering analysis*

As mentioned in the paper, the number of the clusters is chosen based on the Calinski-Harabasz criterion and $k = 4$ is chosen as the number of clusters since its Calinski-Harabasz value is maximal (Figure S6b). In comparison, if we run $k$-mean clustering on the PCA embedding, we also get clusters as shown below (Figure S6a). However, the Calinski-Harabasz criterion (Figure S6b), does not reveal a local maxima as $k$ increases, thus suggesting a lack of clear clustering structure in the low-dimensional embedding of 2sPCA. This validates that compared with our nonlinear embedding, PCA-based linear methods generate less-structured embeddings for the multi-task fMRI data.

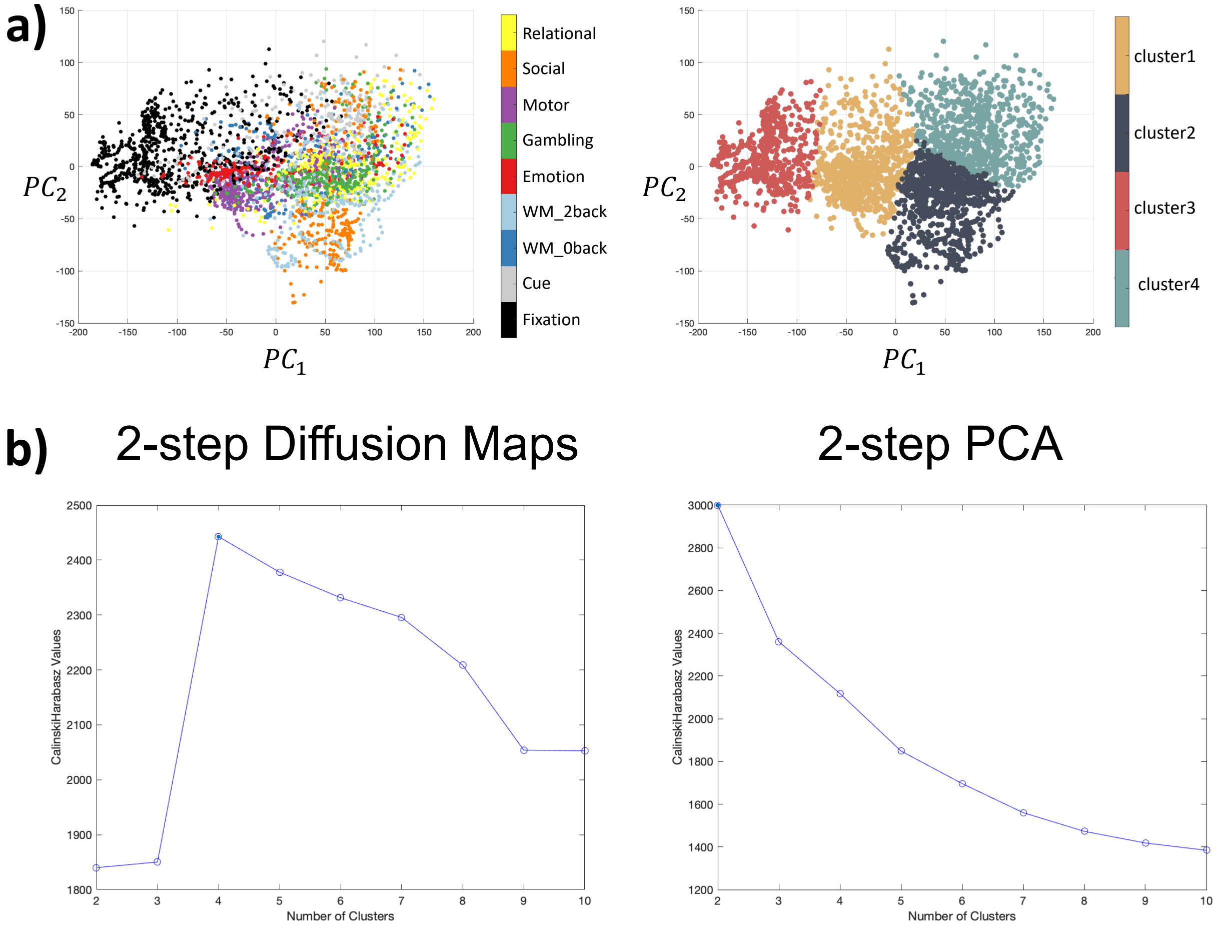


**Figure S6 Brain state clustering comparison.** ***a)*** *K*-means clustering result based on the 2-step PCA embedding. **b)** Clustering evaluation based on the Calinski-Harabasz criterion.

*2sPCA embedding also reveals global integration and segregation*

In the paper, we have shown that 2sDM embedding's first 2 coordinates are highly correlated with the participation coefficient ($B_T$), suggesting that the task embedding is able to capture handcrafted features like $B_T$ in an unsupervised manner (Figure 4). It is also interesting that the 2sPCA embedding also reveals a similar relationship (Figure S7), suggesting that the embedding is also related with the global segregation and integration pattern. Thus, both 2sDM and 2sPCA's embedding is related with $B_T$.

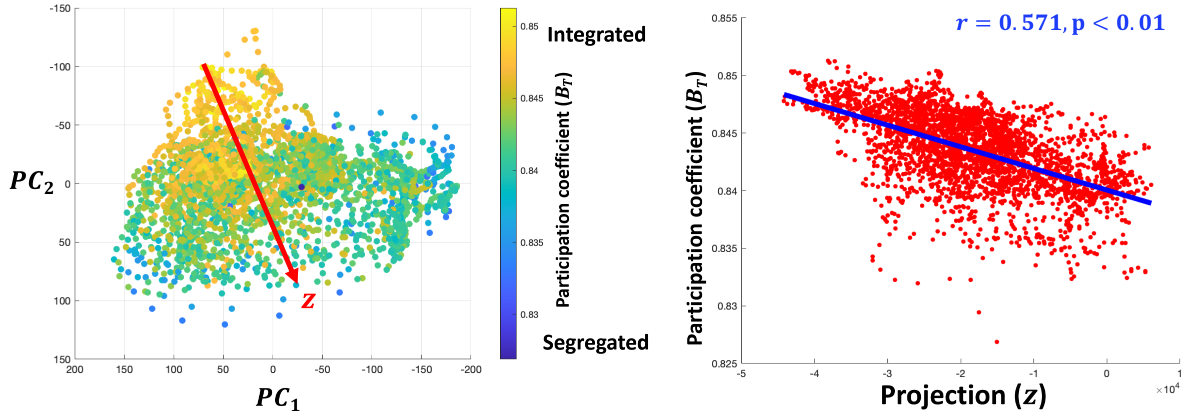


**Figure S7 2sPCA embedding is related with global integration and segregation.** a) 2sDM embedding in HCP dataset colored by the time-resolved $B_T$. b) Scatter plot of the $B_T$ with the projection onto the diagonal of the embedding structure (z). Correlation of $z$ with $B_T$ is shown with a line of best fit. Projection direction $z$ is determined manually to approximate the diagonal direction of the 2-dimensional embedding.

### Nyström extension

Nyström extension is a non-parametric out-of-sample function extension method used to extend the embedding learned with training dataset to unseen data points. Suppose we have a kernel function $K(a, b)$ that generated a symmetric matrix $M$ with entries $M_{ij} = K(x_i, x_j)$ upon a training dataset $D = \{x_1, \ldots, x_n\}$. Let $(v_k, \lambda_k)$ be an (eigenvector, eigenvalue) pair that solves $Mv_k = \lambda_k v_k$. The $k$-th coordinate of diffusion maps embedding is then $\lambda_k v_k$. Let $y_k(x)$ denote the $k$-th diffusion maps embedding associated with a new point $x$, then $y_k(x) = \sum_{i=1}^{n} v_{ki} K(x, x_i)$.

### Low-dimensional embedding in patients with schizophrenia

In addition, to replicate the dimensionality reduction result using all healthy comparison (HC) subjects, we also perform 2sDM on the schizophrenia (SZ) data in the UCLA Consortium for Neuropsychiatric Phenomics (CNP; Poldrack et al., 2016) dataset and produce a similarly shaped embedding as is seen in the HC data (Figure S8), suggesting that similar brain states are present in both HC and SZ. This lays a good foundation for the downstream brain dynamics analyses that are based on brain states, as similar brain states could be identified in both groups. Comparisons between groups could then be made by comparing state-related properties like brain state dwell-time distribution and transition probability. The advantage of our framework compared with existing approaches is that we start from a group-average embedding so that no additional averaging will be needed to infer cohort-wise statistics. Moreover, similar trajectories-based analysis can also be made to compare the cognitive dynamic processes between the two cohorts.

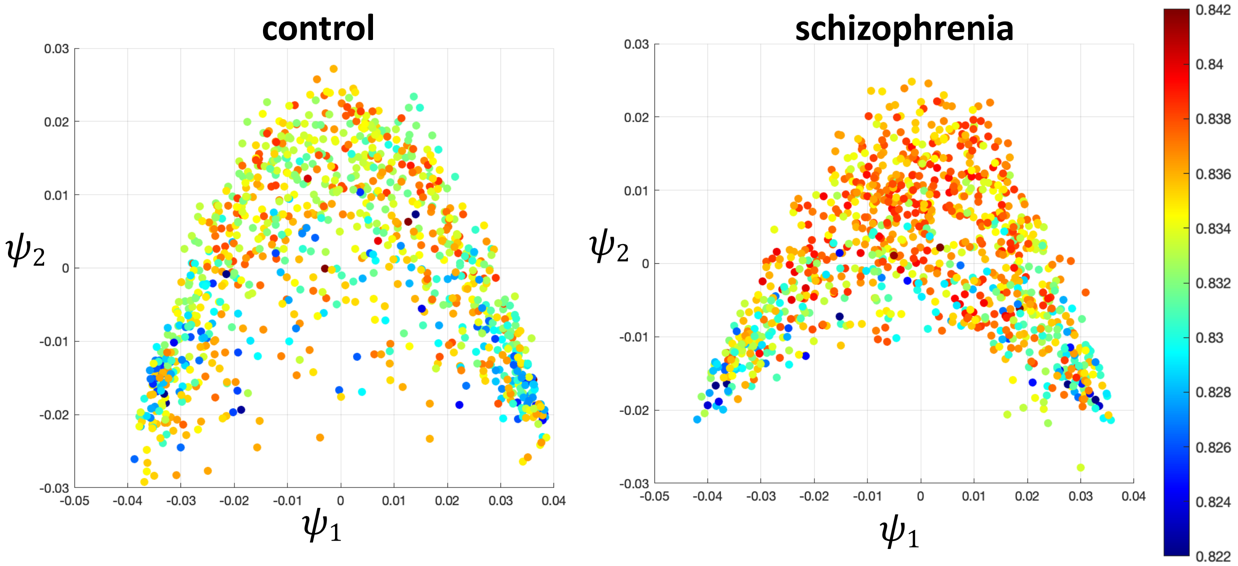


**Figure S8** Embedding with the first 2 dimensions of 2sDM on the HC and SZ groups in CNP dataset, colored by the corresponding participation coefficient ($B_T$) with the same colormap.
